# Identification of plasma proteins associated with oesophageal cancer chemotherapeutic treatment outcomes using SWATH-MS

**DOI:** 10.1101/2022.04.07.487448

**Authors:** Naici Guo, Giorgos Minas, Silvia A. Synowsky, Margaret R. Dunne, Hasnain Ahmed, Rhiannon McShane, Anshul Bhardwaj, Noel E. Donlon, Cliona Lorton, Jacintha O’Sullivan, John V. Reynolds, Peter D. Caie, Sally L. Shirran, Andy G. Lynch, Alan J. Stewart, Swati Arya

## Abstract

Oesophageal adenocarcinoma (OAC) is an aggressive cancer with a five-year survival of <15%. Current chemotherapeutic strategies only benefit a minority (20-30%) of patients and there are no methods available to differentiate between responders and non-responders. We performed quantitative proteomics using Sequential Window Acquisition of all THeoretical fragment-ion spectra-Mass Spectrometry (SWATH-MS) on albumin/IgG-depleted and non-depleted plasma samples from 23 patients with locally advanced OAC prior to treatment. Individuals were grouped based on tumour regression (TRG) score (TRG1/2/3 *vs* TRG4/5) after chemotherapy, and differentially abundant proteins were compared. Protein depletion of highly abundant proteins led to the identification of around twice as many proteins. SWATH-MS revealed significant quantitative differences in the abundance of several proteins between the two groups. These included complement c1q subunit proteins, C1QA, C1QB and C1QC, which were of higher abundance in the low TRG group. Of those that were found to be of higher abundance in the high TRG group, GSTP1 was found to exhibit the lowest p-value and highest classification accuracy and Cohen’s kappa value. Concentrations of these proteins were further examined using ELISA-based assays. This study provides quantitative information relating to differences in the plasma proteome that underpin response to chemotherapeutic treatment in oesophageal cancers.

## Introduction

Oesophageal cancers, including oesophageal adenocarcinoma (OAC) and oesophageal gastric junction cancer (OGJ), are one of the leading causes of cancer mortality worldwide, with a poor 5-year survival rate (globally <20%; developed countries <10%) [1]. It is asymptomatic in early stages and has no biomarkers for its early detection which leads to late diagnosis and poor prognosis resulting in high mortality rate. In 2018, there were an estimated 85,000 new cases of OAC globally (age-standardised incidence rates 0.9 per 100 000, both sexes combined) [2]. The burden of OAC is expected to rise dramatically and surpass oesophageal squamous cell carcinoma, in high-income countries by 2020 [3], with an accompanying rise in the number of patients seeking treatment. Curative therapy consists of surgery, either alone or in combination with adjuvant or neo-adjuvant chemotherapy or radiation, or combination chemoradiotherapy regimens (neo-CRT) [4,5]. However, only a minority of patients respond to such treatments, meaning the majority (70-80%) of patients will experience treatment-related toxicity and delay to surgery with no clinical benefit [6,7]. There are currently no clinico-pathological means of predicting which patients will benefit from chemotherapeutic treatments. There is therefore an urgent need to improve oesophageal cancer disease management and treatment strategies.

Cancer progression and treatment response involve multi-step processes originating from interactions between the tumour and various host cell types and extracellular factors [8]. Moreover, persistent systemic inflammation has been implicated as a possible mechanism of resistance to neo-CRT [9]. Circulating plasma proteins reflect systemic events such as inflammation and angiogenesis and are more amenable to routine interrogation than tumour tissues. Plasma proteins such as serum C-reactive protein (CRP; [10]), soluble interleukin-6 receptor (sIL6R; [11]) and vascular endothelial growth factor (VEGF; [12]) have all demonstrated prognostic potential in oesophageal cancer. In addition to this, there is evidence that plasma proteins can serve as potential prognostic markers for other forms of cancers such as Transforming Growth Factor Beta 1 (TGFβ1) in triple negative breast cancer [13].

Although useful, the previous studies have not comprehensively examined changes in the concentrations of plasma proteins prior to oesophageal cancer treatment. In this study we quantitatively profiled the plasma proteomes of individuals undergoing chemotherapy treatment (mostly MAGIC, but also FLOT or Folfox regimens; [14-16]) using SWATH-MS (Sequential Window Acquisition of All Theoretical Mass Spectra), a data-independent acquisition (DIA)-based method. DIA-based quantitative proteomics represents a powerful tool for identifying the cellular and molecular changes that underlie disease; this is particularly useful in understanding the pathophysiology of cancers and options for their treatment. Here we have used both albumin/IgG-depleted and non-depleted pre-treatment plasma samples to identify differentially abundant proteins in samples grouped according to Mandard tumour regression grade (TRG, with TRG 1 indicating no residual cancer cells, TRG 2 indicating rare residual cells, TRG 3 representing an increase in the number of residual cancer cells, but fibrosis still predominates, TRG 4 where cancer outgrows fibrosis, and TRG 5 represents a complete absence of regressive changes; [17]). This work details proteomic differences that may underpin treatment responses. These in the future could be used to predict response to treatment and allow the development of diagnostic tools to guide personalised treatment options for oesophageal cancer.

## Methods

### Ethical approvals and clinical cohort information

This study was approved by the School of Medicine Ethics Committee, University of St Andrews (Ref: MD9202). Samples were obtained from the Upper Gastrointestinal Trinity Biobank, Trinity Translational Medicine Institute, Trinity College Dublin, St James’ Hospital, Dublin, with approval from Tallaght/St James’s Hospital committee (reference: 2011/27/01). All methods outlined were performed in accordance with the relevant ethical guidelines and regulations. A cohort of 23 patients with locally advanced OAC or OGJ cancers were included for clinical analysis, and these underwent proteomic analysis upon giving informed consent (see Table 1). This cohort received pre- and post-operative chemotherapy as per the MAGIC trial regimen using etoposide, cisplatin or oxaliplatin, and fluorouracil or capecitabine [14], or in a few cases the FLOT regimen consisting of docetaxel, oxaliplatin, leucovirin, and 5-fluorouracil [15] or Folfox regimen consisting of folinic acid 5-fluoruracil and oxaliplatin [16]. The cohort was grouped based on TRG status (TRG1/2/3 vs TRG 4/5).

**Table 1.** Demographic information on the patient cohort by study group.

|  | **TRG 1/2/3**<br>**(n=8)** | **TRG 4 and 5**<br>**(n=15)** |
| --- | --- | --- |
| **Mean age (years)** | 62.9±14.0 | 63.0±8.9 |
| **Sex (no. of males)** | 7 | 15 |
| **Cancer type (no. of individuals)** |  |  |
| Oesophageal adenocarcinoma | 6 | 13 |
| Gastro-oesophageal junction cancer | 2 | 2 |
| **Neo-adjuvant treatment (no. of individuals)** |  |  |
| MAGIC | 6 | 13 |
| FLOT | 2 | 1 |
| Folfox | 0 | 1 |
| **TRG Score (no. of individuals)** |  |  |
| =1 | 3 | 0 |
| =2 | 4 | 0 |
| =3 | 1 | 0 |
| =4 | 0 | 9 |
| =5 | 0 | 6 |
| **Lymphovascular invasion (no. of individuals)** | 1 | 11 |
| **Disease recurrence (no. of individuals)** | 1 | 12 |

### Experimental design and statistical rationale for SWATH-MS

This experiment aimed to identify proteomic differences in plasma that may underlie (or potentially predict) chemotherapy treatment outcomes in OAC patients. Plasma was prepared from whole blood collected in EDTA Vacutainer tubes (BD, Wokingham, UK) taken after diagnosis but prior to chemotherapy. Both non-depleted and albumin/IgG-depleted plasma samples were used. Albumin/IgG depletion was performed according to manufacturer’s instructions using Albumin/IgG depletion spin trap columns (Cytiva, Marlborough, MA, USA). These were analysed independently and grouped based upon the TRG score (grades 1/2/3 *vs* grades 4/5) after treatment. It was not possible to group samples based on post-treatment survival time as detailed data for this parameter was not available for all samples. A sample-specific library was made by pooling all samples for a comprehensive representation of the cohort.

### Sample preparation for mass spectrometry

Plasma containing 30 μg of total protein was digested for mass spectrometric analysis. For this, plasma proteins were denatured in a final concentration of 5 M urea, 2% sodium deoxycholate and 50 mM ammonium bicarbonate. Proteins were reduced and alkylated with 5 mM tris (2-carboxyethyl) phosphine followed by 5 mM iodoacetamide. The reaction was quenched with 10 mM dithiothreitol. Samples were diluted with 50 mM ammonium bicarbonate to a final urea concentration of 1.5 M. The resulting samples were then digested with trypsin (0.2 µg/μl trypsin, 1:50 ratio (w/w); Promega, Southampton, UK) overnight at 30°C. Finally, 0.5% (v/v) trifluoroacetic acid (TFA) was added. Peptides were desalted using a C18 SepPak cartridge (Waters, Elstree, UK). An aliquot of the peptides in each sample was kept separate for the SWATH analysis. The remaining peptides were pooled for a spectral library generation. The solvent in each sample was removed using a SpeedVac (Thermo Fisher Scientific, Loughborough, UK).

### LC-ESI-MSMS analysis for spectral library generation

Peptides used for the library generation were separated using high pH reversed phase fractionation. The pooled, dried peptides were then resuspended in 100 µl Buffer A, consisting of 10 mM ammonium formate, 2% MeCN, pH 10.0. Peptides were then fractionated on a XBridge C18 column (4.6 × 100 mm, 5 µm, Waters) at 1 ml/min with the following gradient: linear gradient of 4-28% Buffer B (10 mM ammonium formate, 90% MeCN, pH 10.0) for 36 min, then 28-50% B for 8 min, followed by 100% B for a further 5 min to wash the column, before re-equilibration in 100% A for 10 min. Fractions of 0.5 ml were collected every 30 s. The UV chromatogram was inspected and fractions pooled to give 25 fractions across the elution profile. The pooled fractions were dried and resuspended in 0.1% FA for mass spectrometric analysis.

The fractions were analysed individually on a Sciex TripleTOF 5600+ system mass spectrometer (Sciex, Framingham, MA, USA) coupled to an Eksigent nanoLC AS-2/2Dplus system, in data dependent mode, to achieve in depth identification of proteins. Additionally, 1 μg of peptides from each individually digested sample for the SWATH analysis were combined and analysed in data dependent mode. Prior to mass spectrometric analysis, reference iRT peptides (Biognosys, Schlieren, Switzerland) were added to each sample according to the manufacturer’s specifications to allow correction of retention times. The samples were loaded in loading buffer (2% MeCN, 0.05% trifluoroacetic acid) and bound to an Acclaim Pepmap 100 µm × 2 cm trap (Thermo Fisher Scientific), and washed for 10 min to waste, after which the trap was turned in-line with the analytical column (Acclaim Pepmap RSLC 75 µm × 15 cm). The analytical solvent system consisted of Buffer A (2% MeCN, 0.1% FA in water) and Buffer B (2% water, 0.1% FA in MeCN) at a flow rate of 300 nl/min, with the following gradient: linear 1-20% of Buffer B over 90 min, linear 20-40% of Buffer B over 30 min, linear 40-99% of Buffer B over 10 min, isocratic 99% of Buffer B for 5 min, linear 99-1% of buffer B over 2.5 min and isocratic 1% solvent buffer B for 12.5 min. The mass spectrometer was operated in data-dependent analysis (DDA) top 20 positive ion mode, with 250 and 150 ms acquisition time for the MS1 (m/z 400-1200) and MS2 (m/z 230-1800) scans respectively, and 15 s dynamic exclusion. Rolling collision energy with a collision energy spread of 5 eV was used for fragmentation. The data was searched against SwissProt restricted to human proteins (accessed January 2019, 42407 proteins) using the Mascot Algorithm with the following search parameters: trypsin as the cleavage enzyme (/K-\P and /R-\P) and carbamidomethylation as a fixed modification of cysteines and oxidation as variable modification. Peptide mass tolerance was set to 20 ppm with fragment mass tolerance set to 0.1 Da. Note that the iRT peptides were included in this database.

### SWATH-MS data acquisition

For SWATH-MS data acquisition, the same mass spectrometer and LC-MS/MS setup was used essentially as described above but operated in SWATH mode. The method uses 100 windows of variable Da effective isolation width with a 1 Da overlap as specified by Sciex. Each window has a dwell time of 150 ms to cover the mass range of 400-1250 m/z in TOF-MS mode and MS/MS data is acquired over a range of 230-1800 m/z with high sensitivity setting and a dwell time of 35 ms, resulting in a cycle time of 3.7 s. The collision energy for each window was set using the collision energy of a 2+ ion centred in the middle of the window with a spread of 5 eV.

### Data processing and availability

Identified proteins within Mascot were imported into Skyline 21.10.146 for spectral library generation. SWATH-MS results were mapped against this library. Skyline parameters were chosen as follows: Peptide settings (trypsin digestion with 1 missed cleavage, human as background proteome, carbamidomethylation (C) and oxidation (M) as modifications); Transition settings (2^+^ and 3^+^ precursor charges, 1^+^ and 2^+^ ion charges, b and y ions and precursor. Six transitions were selected per peptide. The peak area was normalized based on the median value. Moderated t-tests with an empirical Bayes estimate of the variance were performed in R (version 4.1.1) with the limma (version 3.48.3) package [18]. The implementation of the Linear discriminant analysis (LDA) classifier with repeated stratified K-fold cross-validation was implemented with a scikit-learn python package (version 0.24.1) [19]. The mass spectrometry proteomics data have been deposited to the ProteomeXchange Consortium via the PRIDE Partner Repository [20], with the dataset identifier PXD032137.

### Enzyme-linked immunosorbent assays (ELISAs)

ELISAs were performed on the patient plasma samples using commercially available kits for human complement component C1q and human glutathione S-transferase pi-1 (GSTP1; Cloud-Clone Corp., Buckingham, UK). Samples were centrifuged at 13,000 × g for 10 min and the supernatant used for analysis. The assay was performed in accordance with the manufacturer’s instructions.

## Results

### Comparison of proteomic data from depleted vs non-depleted plasma samples taken from patients prior to undergoing chemotherapy for OAC

For SWATH-MS quantitation, cohort-specific reference spectral libraries were generated for both depleted and non-depleted sample sets (for use as a reference in generating peptide query parameters for the peptide centric analysis of the data) by data-dependent acquisition analysis of the plasma proteins. To determine the differences in levels of peptides/proteins present across the different TRG groups, we measured quantitative proteome profiles using albumin/IgG-depleted and non-depleted plasma for each individual sample by SWATH-MS. The resultant data were log_2_ transformed and then normalised using median normalization at the protein and peptide level for non-depleted data and depleted data, respectively. As shown in Figures S1, S2, normalisation effectively scaled the patient samples to the same median and stabilised variance across the mean abundance of proteins and peptides. Comparison of the obtained quantitative data identified 1,640 and 2,870 peptides (1% FDR) in non-depleted and depleted samples, respectively. These represented 330 proteins in non-depleted samples and 537 proteins in depleted samples. There was a significant overlap between the two datasets where 1529 peptides (from 295 proteins) were consistently found in both non-depleted and depleted data (Figure 1A). The Bland-Altman analysis was performed for estimating the degree of concordance between the measurement of relative protein abundance level for commonly found 295 proteins. As shown in Figure 1B, the mean difference is –2.7, which implies, on average depleted samples have 2.7 units higher in relative protein abundance than non-depleted samples. Most of the proteins lie in 95% limits of agreement, which are [–6.85, 1.45], except for outlier proteins that are likely to have been affected by depletion, as shown in Table S1. Comparing the distributions of the means of proteins between non-depleted and depleted data (Figure 1C), consistently found proteins showed higher mean in depleted data than non-depleted data. The correlation of relative protein abundance for consistently found proteins between non-depleted and depleted samples is shown in Figure 1D (Pearson correlation coefficient = 0.84). Following both protein and peptide normalization steps, serum albumin and immunoglobin proteins/peptides were relatively more abundant in the non-depleted data than in the depleted data, as expected.

**Figure 1.**
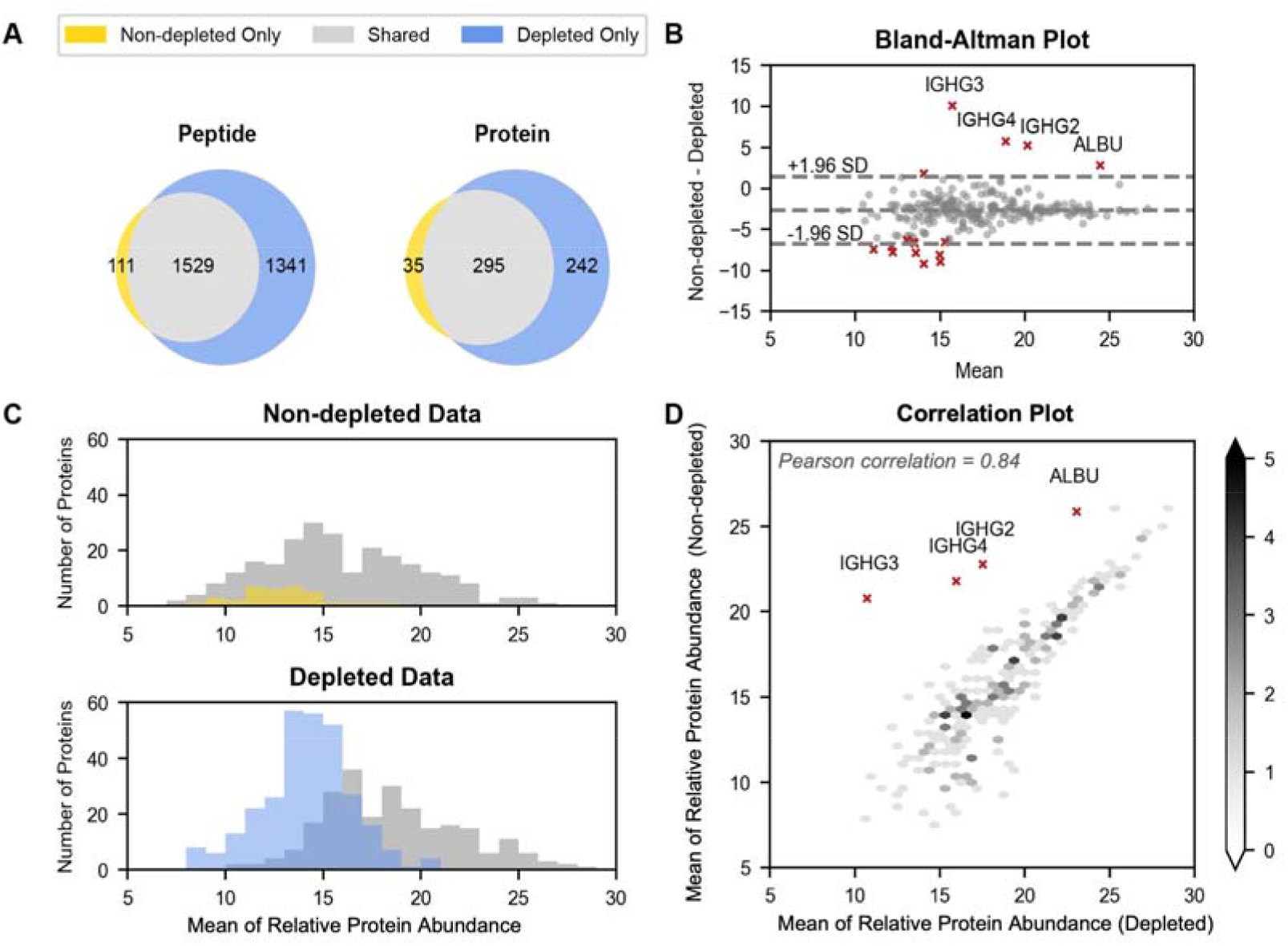
Comparison of proteomic data from depleted *vs*. non-depleted plasma samples. Grey represents peptides or proteins that are detected in both non-depleted data and depleted data. Yellow and blue indicate peptides or proteins that are uniquely existing in non-depleted data and depleted data, respectively. A. Venn diagrams of 1640 and 2870 peptides, 330 and 537 proteins in non-depleted data and depleted data, respectively. B. Bland-Altman Plot showing agreement between depleted and non-depleted data (log_2_ relative abundance) for commonly found proteins. C. Overlay histograms of means of commonly found proteins from non-depleted data and depleted data. D. Scatterplot coloured by density shows the correlation between depleted and non-depleted data for all common proteins. Note that albumin and immunoglobins have a higher abundance in non-depleted data than in depleted data as expected.

### Quantitative analysis of proteins from plasma taken from OAC patients prior to undergoing chemotherapy

#### Univariate analysis

Relative protein abundance in non-depleted plasma samples was compared between low and high TRG groups (Figure S4). Of the proteins consistently quantified, the relative abundance of a total of 14 unique proteins were statistically significantly altered (p-value <0.05) between TRG1/2/3 and TRG4/5 groups. The quantitative data for all differentially abundant proteins is presented. The proteins HBB, C1RL and FCGBP, where quantitation was based on more than one peptide, exhibited differences in relative abundance between the two groups. Relative protein abundance in depleted plasma samples was also compared between low and high TRG groups. A volcano plot showing a comparison of the relative abundance of plasma proteins between TRG1/2/3 and TRG4/5 is presented in Figure 2A. Of the proteins consistently quantified, the relative abundance of a total of 24 unique proteins (12 based on data from more than one peptide) were statistically significantly altered (p-value <0.05) between TRG1/2/3 and TRG4/5 groups. Adjusted p-values (q-values) were computed to control the False Discovery Rate (FDR) using the Benjamini-Hochberg method. These values are in line with results of similar sample size in proteomics and not used in determining significance. The quantitative data for all differentially abundant proteins is presented in Figure 2B. Box-and-whisker plots showing the relative abundance of differentially abundant proteins in the two groups where data is based on more than one peptide (GSTP1, C1QC, BGH3, C1QB, CO2, FBN1, VASN, C4BPA, CYTM, LSAMP, CRP and CALL5), is shown in Figure 2C.

**Figure 2.**
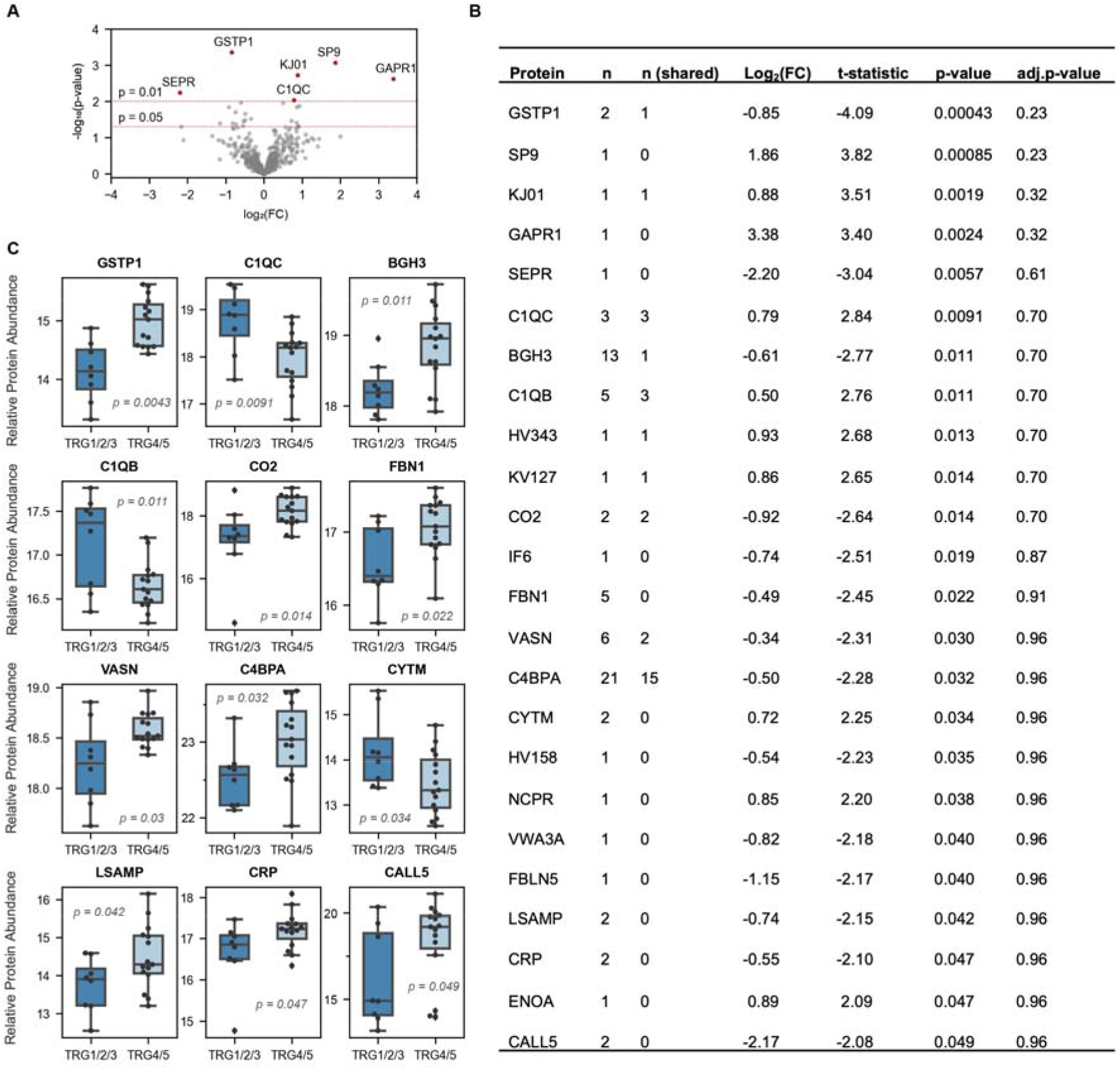
Relative abundance of proteins in depleted plasma based on the univariate analysis comparing samples from individuals with low (1/2/3) and high (4/5) TRG scores. A. Volcano plot showing log_2_-fold change (x-axis) versus negative log_10_ of p-value (y-axis). Red highlighted dots represent proteins with p-value <0.01 identified by the empirical Bayes test. B. Quantitative data for proteins with p-value < 0.05. The columns represent the following: n = number of peptides identified from depleted plasma sample; n (shared) = number of peptides that are commonly found in non-depleted plasma samples). C. Boxplots showing the relative expressions of differentially expressed proteins between TRG1/2/3 and TRG4/5 groups. Only those proteins where quantification is based on more than one peptide are shown.

#### Multivariate analysis

In addition to the univariate analysis, the LDA classifier that predicts sample class based on multiple predictor variables is applied. To fit this classifier, relative peptide abundance that are from the same protein are considered as predictor variables and TRG1/2/3 and TRG4/5 as two sample classes. Cross-validation was used to test the ability of the LDA classifier to make predictions on data that are not used for fitting the model. Thus, for each protein, an LDA model is fitted with a repeated stratified K-fold cross-validator to examine the model performance. Patient samples are randomly partitioned into K (K=4) segments. K-1 segments (75% of patients) are used for training the LDA model and the remaining one segment (25% of patients) is used as a held-out segment for testing the predictions. The testing here is a comparison of predictions with actual TRG groups. The fraction of correctly classified samples is recorded as a classification accuracy score, e.g., a classification accuracy score of 0.8 implies that 80 % of patients in the held-out segment are allocated correctly. The partitioning into K groups of samples is done using a stratified sampler to ensure the proportional representation of the low and high TRG patients in the K segments. After K iterations, each K segment has been held out for testing, thus K classification accuracy scores are generated for each cross-validator. This K-fold cross-validation by stratified-random partitioning is repeated 100 times. The accuracy score presented in Figure S4A and Figure 3A represents the average of 400 classification accuracy scores that are from 100 repeats of four held-out testing sets. Cohen’s kappa statistics, which compares the accuracy of LDA with an accuracy of purely random guessing is also computed [21]. The average kappa statistics and confidence interval are calculated to measure the consistency of predictions in 400 held out sets, ranging from –1(total disagreement) through 0 (random classification to 1(total agreement). Three proteins exhibited an average Cohen’s kappa value greater than 0.35 in non-depleted plasma samples; these were IGI, APOD and TETN. Box-and-whisker plots showing the relative abundance of these three proteins is shown in Figure S5B. Multivariate analysis of the depleted plasma peptide data was also performed using the LDA model. Fourteen proteins exhibited an average kappa value greater than 0.35. These were GSTP1, GAPR1, KJ01, SP9, C1QC, A1AT, SEPR, SEPP1, C1QA, KV127, ENOA, HV108, ACPH and C1QB (Figure 3A). Box-and-whisker plots showing the relative abundance of proteins, where data is based on more than one peptide is shown in Figure 3B. Interestingly, nine proteins displayed both significant differential abundance at the protein level and high accuracy at the peptide level when comparing the two groups. These include GSTP1, GAPR1, KJ01, SP9, C1QC, SEPR, KV127, ENOA and C1QB.

**Figure 3.**
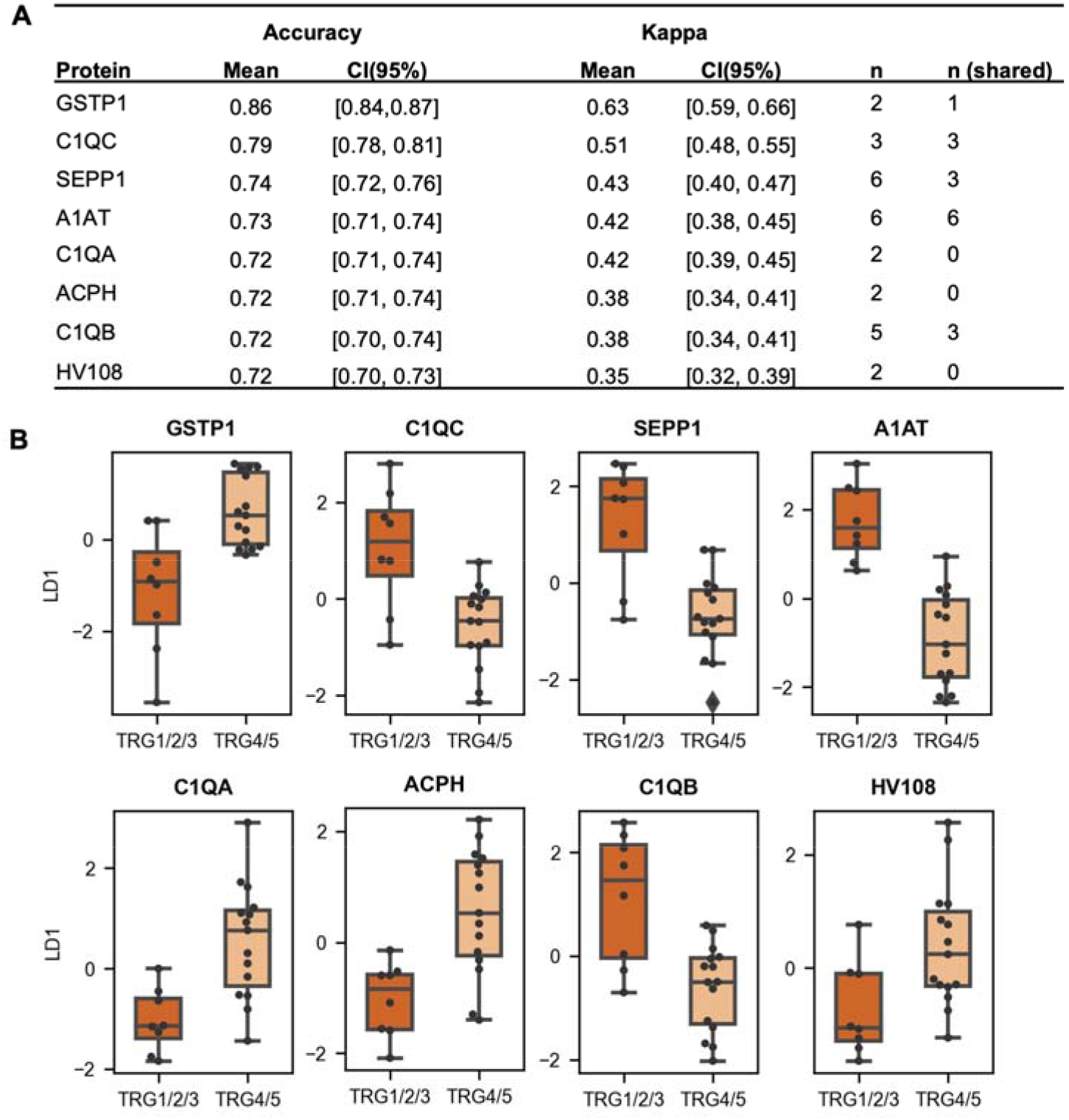
Prediction of low or high TRG score groups from proteins with multiple peptides in the depleted data. A linear discriminant analysis (LDA) classifier is applied to all peptides from the protein. A. Table presents the results of the LDA classifier for the proteins that have the highest average Cohen’s kappa. The proteins are ranked by the average Cohen’s kappa and the top hits are listed. The columns represent the following: CI= confidence interval; n = the number of peptides identified from depleted plasma sample; n (shared) = the number of peptides that are commonly found in non-depleted plasma samples). B. Linear discriminant scores regarding low (1/2/3) and high (4/5) TRG scores groups. Significant differences along the first linear discriminant LD1 were detected for all the selected proteins, suggesting that low and high TRG score groups can be very well separated by the weighted sum of relative peptide expression level from each protein, as shown in Figure S6.

### Functional annotation of differentially expressed proteins

Pathway enrichment analysis of plasma proteins (depleted and non-depleted) displaying differential abundance between TRG 1/2/3 vs TRG 4/5 groups was performed using Metascape (https://metascape.org; [22]). This is an over-representation analysis that statistically determines whether proteins from certain pathways are enriched in the data. Top clusters along with their representative enriched terms were selected. Resultant regulated pathways in TRG 1/2/3 group are shown in Figure 4A and those that are regulated in TRG 4/5 are shown in Figure 4B. The results show that the proteins that are expressed at higher levels in patients under TRG 1/2/3 group are enriched in biological processes related to “C1Q complex”, “Platelet degranulation”, “Selenium micronutrient network”, “Negative regulation of endopeptidase activity” and “Monocarboxylic acid metabolic process”. Proteins that are of higher abundance in patients within the TRG4/5 group are involved in processes such as “Regulation of superoxide metabolic process”, “Naba extracellular matrix glycoproteins”, “Complement cascade”, “Angiogenesis”, “Negative regulation of Immune system”, “Hemostasis”, “Negative regulation of cell differentiation” and “Response to hormone”.

**Figure 4.**
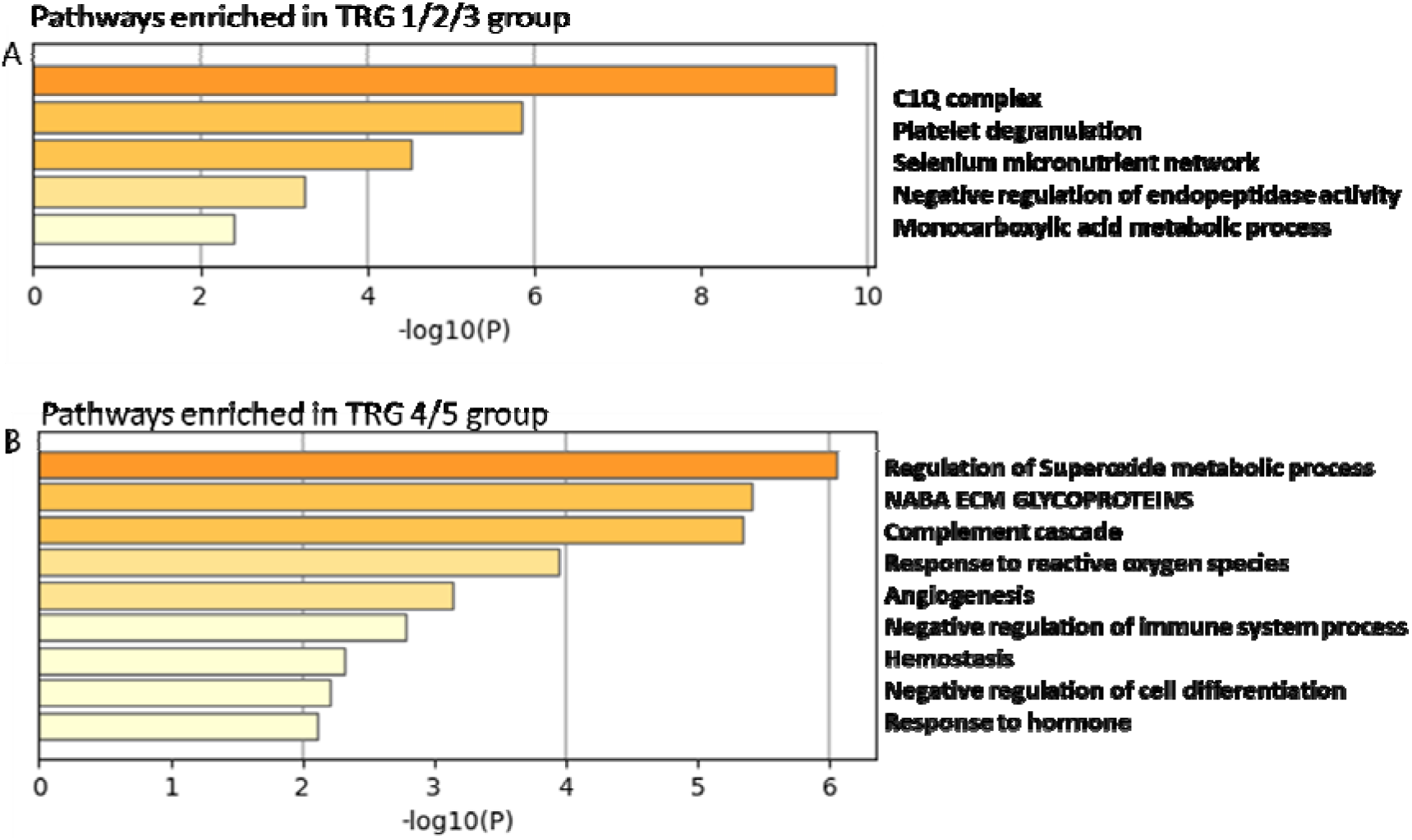
Enriched Ontology Clusters of significantly altered proteins. Enriched pathway clusters developed from all plasma proteins that display significantly altered expression levels between individuals with low (1/2/3) and high (4/5) TRG scores. A. Pathways related to all upregulated proteins in individuals with TRG1/2/3 and B. Pathways related to all upregulated proteins in individuals with TRG 4/5.

### Determination of plasma concentrations of complement component C1q complex in patient samples

Of the proteins that exhibited differing mean relative abundances between groups, C1q (complex) and GSTP1 were chosen for further analysis. To validate the findings from the SWATH-MS analysis, ELISA assays were carried out on the non-depleted plasma samples. The resultant data are shown in Figure 5. Statistically significant differences in the concentrations of C1q complex and GSTP1 were observed in agreement with the SWATH-MS data. Moreover, the predictive performance of c1q complex was examined and compared with C1QA, C1QB and C1QC subunits. As illustrated in Figure 5B, greater separation along the first linear discriminant was observed when using relative abundance of all the peptides from c1q protein complex as features.

**Figure 5.**
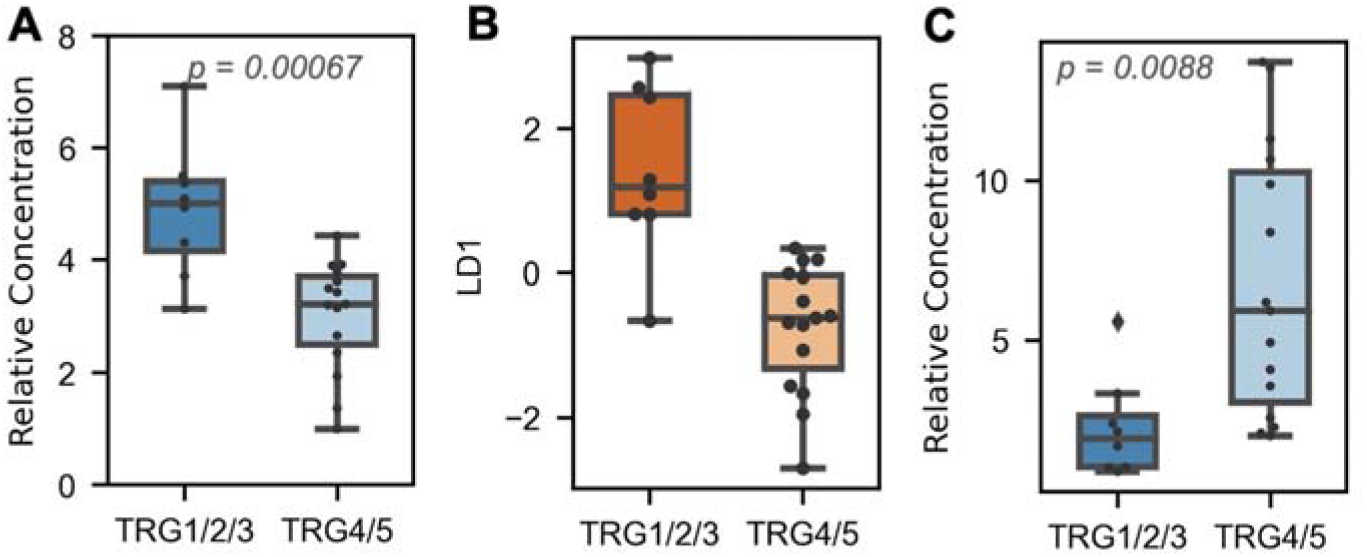
Quantitative analysis of C1q and GSTP1 in plasma samples by enzyme-linked immunosorbent assay (ELISA) and comparison of C1q protein complex and subunit proteins in respect of predicting TRG score groups. Statistical comparisons were carried out by an independent two-sample t-test. A. Relative concentrations of C1q in samples associated with a low (1/2/3) vs high TRG score (4/5). B. Linear discriminant scores regarding low (1/2/3) and high (4/5) TRG scores for C1q protein complex. Compared with individual C1q subunit proteins, greater difference was observed along LD1 axis, indicating improved classification performance based on weighted sum of relative peptide abundance level from c1q complex. C. Relative concentrations of GSTP1 in samples associated with a low (1/2/3) vs high TRG score (4/5). C. Relative concentrations of GSTP1 in samples associated with a low (1/2/3) vs high TRG score (4/5).

## Discussion

SWATH-MS was employed to identify proteomic differences in plasma taken from OAC patients prior to undergoing chemotherapy. In our study we examined both native plasma and plasma depleted of abundant proteins using a commercial Albumin/IgG-depletion kit. To our knowledge, this is the first SWATH-MS-based clinical study to directly compare results obtained from both depleted and non-depleted plasma samples. Depletion led to the identification and relative quantitation of many more peptides/proteins (∼2-times as many) than could be observed when analysing the non-depleted plasma. A comparison of peptides identified in both depleted and non-depleted samples showed a high level of agreement in the measurement obtained using linear regression, suggesting that although depletion would be expected to alter the variability of some proteins (i.e., those that bound to the column which included albumin, IgG proteins and ATP binding cassette subfamily member 1) it did not significantly alter the relative abundance of the vast majority of proteins in plasma. Thus, our study supports the utilisation of protein depletion as a viable strategy to gain higher peptide/protein coverage in plasma using SWATH-MS or similar approaches.

It was evident from the clinical data that patients who would go on to have a higher TRG score (which equates to a lower degree of tumour regression after chemotherapy) were more likely to have tumours that developed lymphovascular invasion and have recurrence of disease. To examine whether there are any plasma proteomic differences between groups with high and low TRG, the quantitative proteomic data for individuals in each group were compared and differences in several proteins were observed between these groups. Univariate analysis of the non-depleted data revealed 14 plasma proteins that had significantly different abundance between those with a low TRG compared to a high TRG (p-value <0.05). However, it is important to add that the results of 11 of these were based on only one detected peptide. Multivariate analysis of proteins (where more than one peptide was detected) was also performed but did not identify any proteins with high classification accuracy in the non-depleted dataset.

With the albumin/IgG-depleted sample set, we obtained much richer data characterised by increased peptide/protein coverage and a higher signal-to-noise ratio, which allowed the identification of differentially abundant proteins between low and high TRG groups using multivariate analysis. Proteins that exhibited differential abundance between groups using this method included GSTP1, SEPR, BGH3, CO2 and IF6, which were present at a lower concentration, and C1QA, C1QB, C1QC, SP9, GAPR1, HV343 and KV127, which were present at a higher concentration in patients with a lower TRG score after chemotherapy. Among these, the three C1q proteins (C1QA, C1QB and C1QC) were perhaps the most notable as they together form the functional C1q complex. Complement factor C1q, together with C1r and C1s form the C1 complex, the first component in the classical complement pathway. C1q is composed of six identical subunits, each of which is composed of three different polypeptide chains, C1qA, C1qB and C1qC [23]. All three C1q component proteins were more abundant in the plasma of individuals with low TRG scores compared to high TRG scores, suggesting it may play a role in reducing oesophageal cancer severity and could serve as a positive prognostic indicator for this cancer type. Using an ELISA-based assay, the levels of C1q complex were also found to be higher in samples from those who would go on to develop a lower TRG score after chemotherapy. Activation of the classical pathway occurs when C1q binds to an antigen-antibody complex [23]. However, C1q can also exert functions unrelated to complement activation. In cancer, C1q is expressed in the stroma and vascular endothelium of several human malignant tumours [24]. Other proteomic studies have identified C1q as a potential prognostic indicator. In lung and kidney cancer and in glioma, higher C1q levels are associated with a negative prognosis [25,26]. Conversely, in breast cancer and some types of lung cancer, as with our study, higher C1q levels are associated with a more positive prognosis [27,28]. C1q is an important modulator of inflammation and cytokine secretion. What is not clear is whether (and when) C1q is protective against or supportive of cancer progression [27]. C1q can be detrimental to cancer cell viability via cell lytic mechanisms [29,30]. However, C1q can also exert tumour-promoting functions independent of classical pathway activation, as observed for some types of cancers [23,31,32].

Glutathione S-transferase pi-1 (GSTP1) was identified from our multivariate analysis as the protein exhibiting most significantly different abundance in plasma between groups (highest accuracy and Cohen’s kappa value). This finding was further corroborated by ELISA using non-depleted plasma samples. Higher GSTP1 levels were found in individuals with a high TRG score and was thus associated with a poorer prognosis from those in our study. GSTP1 is a member of a family of enzymes that play an important role in detoxification by catalysing the conjugation of hydrophobic and electrophilic compounds with reduced glutathione [33]. *GSTP1* is a polymorphic gene encoding active, functionally different GSTP1 variant proteins that function in xenobiotic metabolism and play a role in susceptibility to cancer, and other diseases [33]. Various members of the glutathione S-transferase (GST) family have been reported as being overexpressed in several cancers and in most cases have been linked to poor prognosis and chemoresistance [34-37]. GSTP1 is involved in cell maintenance, cell survival and the cellular stress response via the NF-κB and MAP kinase pathways during tumour progression. Furthermore, *GSTP1* knockdown in cervical cancer cells showed that evasion of apoptosis was affected, and tumour proliferation was significantly reduced [38]. Further work is required to fully understand the role of GSTP1 and its prognostic relevance in OAC and other cancer types.

Among the other proteins that displayed altered abundance between the two groups based on the univariate analysis, BGH3, CO2, C4BPA, LSAMP, CALL5, CRP, FBN1, and VASN were found to be of higher abundance on average in plasma from patients with a high TRG score after treatment. BGH3 (transforming growth factor-beta-induced protein ig-h3) is thought to play a role in angiogenesis which is important for tumour growth and expansion [39]. A microarray study conducted by He et al. identified significant up-regulation of BGH3 in oesophageal squamous cell carcinoma (OSCC) [40]. CO2 (cytochrome c oxidase subunit 2) is predicted to be involved in regulation of mitochondrial electron transport and has been implicated in cancer progression [41]. C4BP (complement component 4 binding protein) is a glycoprotein responsible for regulating the classical pathway of the complement system, which exists as distinct isoforms made up of alpha and beta subunits; C4BPA, which exhibited differential plasma abundance between groups in our study, encodes the alpha chain [42]. A recent study by Sogawa et al. demonstrated that the sera levels of C4BPA was significantly higher in the preoperative Pancreatic ductal adenocarcinoma (PDAC) patients than in the postoperative ones, C4BPA was also significantly higher in PDAC patients in comparison to healthy controls [43]. LSAMP (limbic system-associated membrane protein) has been reported to be a candidate tumour suppressor gene in a range of cancer types [44-48]. Interestingly, the downregulation of LSAMP expression in cancer cell lines is associated with poor survival prognosis by promoting lung cancer progression [49]. CALL5 (calmodulin-like protein 5) is also thought to have tumour suppressor activities and is silenced at an early stage of carcinogenesis in squamous cell carcinoma of uterine cervix [50]. However, in our study a higher abundance of LSAMP and CALL5 in plasma were associated with a relatively poorer prognosis for oesophageal cancer. How plasma LSAMP and CALL5 levels relate to the cellular expression/activities of these proteins is unknown but it is possible that they may not correlate positively. CRP (C-reactive protein) is a classical inflammatory marker identified to be related to progression of oesophageal cancer and, consistent with our study, elevated pre-treatment serum CRP levels are associated with poorer prognosis in oesophageal cancer patients [51]. FBN1 (fibrillin-1) is an extracellular matrix glycoprotein that provides structural and regulatory support to both elastic and non-elastic connective tissues via calcium-binding microfibrils [52]. This protein has previously been associated with poor survival in ovarian cancer [53] and been proposed as a germ cell tumor marker [54]. VASN (Vasorin/SLITL2) is a classic type 1 transmembrane protein which is linked with vascular injury repair and is overexpressed in a range of tumours [55]. Serum vasorin levels are upregulated in patients with colon cancer, with levels further increasing as cancer progresses [56]. In addition to the C1q subunit proteins, univariate analysis revealed one other protein found to be present at higher abundance in plasma from patients with a low TRG score after treatment, CYTM (Cystatin M). Several studies have found this protein to have anti-tumour activity and to be a positive prognostic indicator for treatment of various cancers [57-59].

From the multivariate analysis of proteomic data, proteins that displayed altered abundance between the two groups (other than GSTP1 and C1q subunit proteins) included ACPH and HV108, which were found to be of lower abundance in those with a low TRG score after treatment, and A1AT and SEPP1 which were more abundant in plasma from patients in this group. ACPH (acyl carrier protein phosphodiesterase) and HV108 (immunoglobulin heavy-chain variable 1-8 protein) have not, to our knowledge, previously been linked to cancer progression or prognosis. A1AT (alpha-1 antitrypsin) is a circulating glycoprotein that inhibits neutrophil elastase and other serine proteases in blood and tissues. The role of this protein in cancer progression is controversial, while deficiency of A1AT is a risk factor for several cancer types, increased plasma/serum expression has been reported for multiple cancers compared to healthy controls [60]. It has also been reported that A1AT levels in the blood are significantly decreased in non-small cell lung cancer and prostate cancer after chemotherapeutic treatment compared with those before the treatment started [61]. SEPP1 (selenoprotein P) is a protein that is positively associated with cancer risk and can regulate tumorigenesis and progression through its effects on cancer-related signalling pathways [62]. However, a protective effect of selenium supplementation on OAC risk, mediated through the antioxidant activity of selenoenzymes, has been reported [63].

Functional annotation of proteins displaying differential abundance between the groups highlighted several pathways that were significantly represented by these proteins. Unsurprisingly, “C1q complex” was the most significantly enriched pathway, being associated with proteins upregulated in the TRG1/2/3 group. Interestingly, pathways classically associated with cancer progression including angiogenesis and negative regulation of both the immune system and cell differentiation were highlighted as involving upregulated proteins in the TRG4/5 group and thus, were associated with poorer treatment outcomes. Although complement C1q was associated with better treatment outcomes, “Complement cascade” was identified as a functional group for proteins associated with poor treatment outcomes. This was interesting as a previous (serum) proteomic study by our group showed that pretreatment serum C4a and C3a levels were significantly higher in poor responders versus good responders [64].

## Conclusions

In conclusion, this study has utilised a combination of quantitative proteomics and statistical methods to identify proteins that have predictive power to display differential abundance between OAC patients that respond favourably and less favourably to chemotherapy. In our study, we analysed both albumin/IgG-depleted and non-depleted plasma in parallel. It was clear that depletion of very highly abundant proteins led to an improvement in signal-to-noise and the consequent detection and quantitation of many more proteins. A direct comparison between proteins common to both depleted and non-depleted sample sets showed a high degree of agreement. From the proteins that exhibited differential abundance, the c1q family proteins and GSTP1 had the highest potential to predict likelihood of tumour regression in response to neo-adjuvant chemotherapy in OAC. These observations point towards the use of such markers for stratifying patients prior to treatment and the further use of quantitative proteomics in personalised medicine to predict treatment outcomes. This study provides a platform for further work, utilising larger sample sets across different treatment regimens for oesophageal cancer, that will aid the development of prognostic assays for use in the clinic.

## Data availability

The proteomic data are available and have been deposited to the ProteomeXchange Consortium via the PRIDE Partner Repository with the dataset identifier PXD032137.

## Acknowledgements

We wish to thank the Wellcome Trust for funding the purchase of the TripleTOF 5600+ mass spectrometer [grant number: 094476/Z/10/Z] and their Institutional Strategic Support Fund [grant number: 097831/Z/11/Z] for funding a transition fellowship (to S.A.). We also wish to thank Tenovus Scotland for PhD studentship funding [grant number: T19-05]; to R.M and H.A. M.R.D. is funded by the Health Research Board, Ireland [grant number: HRB-ILP-POR-2017-055].

## References

1. GBD 2017 Oesophageal Cancer Collaborators. The global, regional, and national burden of oesophageal cancer and its attributable risk factors in 195 countries and territories, 1990-2017: a systematic analysis for the Global Burden of Disease Study 2017. Lancet Gastroenterol Hepatol. 2020;5:582–597.

2. Arnold M, Ferlay J, van Berge Henegouwen MI, Soerjomataram I. Global burden of oesophageal and gastric cancer by histology and subsite in 2018. Gut 2020;69:1564–71.

3. Arnold M, Laversanne M, Brown LM, Devesa SS, Bray F. Predicting the Future Burden of Esophageal Cancer by Histological Subtype: International Trends in Incidence up to 2030. Am J Gastroenterol. 2017;112:1247–1255.

4. Donlon NE, Davern M, Sheppard A, Power R, O’Connell F, Heeran AB, King R, Hayes C, Bhardwaj A, Phelan JJ, Dunne MR, Ravi N, Donohoe CL, O’Sullivan J, Reynolds JV, Lysaght J. The prognostic value of the lymph node in oesophageal adenocarcinoma; incorporating clinicopathological and immunological profiling. Cancers 2021;13:4005.

5. Geh JI, Bond SJ, Bentzen SM, Glynne-Jones R. Systematic overview of preoperative (neoadjuvant) chemoradiotherapy trials in oesophageal cancer: Evidence of a radiation and chemotherapy dose response. Radiotherapy and Oncology 2006;78:236–44.

6. Kelsen DP, Ginsberg R, Pajak TF, Sheahan DG, Gunderson L, Mortimer J, Estes N, Haller DG, Ajani J, Kocha W, Minsky BD, Roth JA. Chemotherapy followed by surgery compared with surgery alone for localized esophageal cancer. New England Journal of Medicine 1998;339:1979–84.

7. Korst RJ, Kansler Al, Port JL, Lee PC, Kerem Y, Altorki NK. Downstaging of T or N predicts long-term survival after preoperative chemotherapy and radical resection for esophageal carcinoma. Annals of Thoracic Surgery 2006;82:480–4.

8. Castro-Giner F, Aceto N. Tracking cancer progression: from circulating tumor cells to metastasis, Genome Medicine 2020;12:13.

9. Marcone S, Buckley A, Ryan CJ, McCabe M, Lynam-Lennon N, Matallanas D, O’Sullivan J, Kennedy S. Proteomic signatures of radioresistance: Alteration of inflammation, angiogenesis and metabolism-related factors in radioresistant oesophageal adenocarcinoma. Cancer Treatment and Research Communications 2021;27:100376.

10. Lorton CM, Higgins L, O’Donoghue N, Donohoe C, O’Connell J, Mockler D, Reynolds JV, Walsh D, Lysaght J. C-reactive protein and C-reactive protein-based scores to predict survival in esophageal and junctional adenocarcinoma: Systematic review and meta-analysis. Annals of Surgical Oncology 2022;29:1853–1865.

11. Makuuchi Y, Honda K, Osaka Y, Kato K, Kojima T, Daiko H, Igaki H, Ito Y, Hoshino S, Tachibana S, Watanabe T, Furuta K, Sekine S, Umaki T, Watabe Y, Miura N, Ono M, Tsuchida A, Yamada T. Soluble interleukin-6 receptor is a serum biomarker for the response of esophageal carcinoma to neoadjuvant chemoradiotherapy. Cancer Science 2013;104:1045–51.

12. Kleespies A, Guba M, Jauch KW, Bruns CJ. Vascular endothelial growth factor in esophageal cancer. Journal of Surgical Oncology 2004;87:95–104.

13. Katayama H, Tsou P, Kobayashi M, Capello M, Wang H, Esteva F, Disis ML, Hanash S. A plasma protein derived TGFβ signature is a prognostic indicator in triple negative breast cancer. NPJ Precision Oncology 2019;3:10.

14. Cunningham D, Allum WH, Stenning SP, Thompson JN, Van de Velde CJ, Nicolson M, Scarffe JH, Lofts FJ, Falk SJ, Iveson TJ, Smith DB, Langley RE, Verma M, Weeden S, Chua YJ. MAGIC Trial Participants, Perioperative chemotherapy versus surgery alone for resectable gastro-esophageal cancer. New England Journal of Medicine 2006;355:11–20.

15. Al-Batran SE, Hofheinz RD, Pauligk C, Kopp HG, Haag GM, Luley KB, Meiler J, Homann N, Lorenzen S, Schmalenberg H, Probst S, Koenigsmann M, Egger M, Prasnikar N, Caca K, Trojan J, Martens UM, Block A, Fischbach W, Mahlberg R, Clemens M, Illerhaus G, Zirlik K, Behringer DM, Schmiegel W, Pohl M, Heike M, Ronellenfitsch U, Schuler M, Bechstein WO, Königsrainer A, Gaiser T, Schirmacher P, Hozaeel W, Reichart A, Goetze TO, Sievert M, Jäger E, Mönig S, Tannapfel A. Histopathological regression after neoadjuvant docetaxel, oxaliplatin, fluorouracil, and leucovorin versus epirubicin, cisplatin, and fluorouracil or capecitabine in patients with resectable gastric or gastro-oesophageal junction adenocarcinoma (FLOT4-AIO): results from the phase 2 part of a multicentre, open-label, randomised phase 2/3 trial. Lancet Oncology 2016;17:1697–708.

16. Conroy T, Galais MP, Raoul JL, Bouché O, Gourgou-Bourgade S, Douillard JY, Etienne PL, Boige V, Martel-Lafay I, Michel P, Llacer-Moscardo C, François E, Créhange G, Abdelghani MB, Juzyna B, Bedenne L, Adenis A, Fédération Francophone de Cancérologie Digestive and UNICANCER-GI Group. Definitive chemoradiotherapy with FOLFOX versus fluorouracil and cisplatin in patients with oesophageal cancer (PRODIGE5/ACCORD17): Final results of a randomised, phase 2/3 trial, Lancet Oncology 2014;15:305–14.

17. Mandard A-M, Dalibard F, Mandard J-C, Marnay J, Henry-Amar M, Petiot J-F, Roussel A, Jacob J-H, Segol P, Samama G, Ollivier J-M, Bonvalot S, Gignoux M. Pathologic assessment of tumor regression after preoperative chemoradiotherapy of esophageal carcinoma. Clinicopathologic correlations. Cancer 1994;73:2680–2686.

18. Ritchie ME, Phipson B, Wu D, Hu Y, Law CW, Shi W, Smyth GK. limma powers differential expression analyses for RNA-sequencing and microarray studies. Nucleic Acids Research 2015;43:e47.

19. Pedregosa F, Varoquaux G, Gramfort A, Michel V, Thirion B. Scikit-learn: Machine learning in Python. Journal of Machine Learning Research 2011;12:2825–30.

20. Perez-Riverol Y, Csordas A, Bai J, Bernal-Llinares M, Hewapathirana S, Kundu DJ, Inuganti A, Griss J, Mayer G, Eisenacher M, Perez E, Uszkoreit J, Pfeuffer J, Sachsenberg T, Yilmaz Ş, Tiwary S, Cox J, Audin E, Walzer M, Jarnuczak AF, Ternent T, Brazma A, Vizcaíno JA. The PRIDE database and related tools and resources in 2019: improving support for quantification data. Nucleic Acids Research 2019;47:D442–D450.

21. Cohen JA. Coefficient of Agreement for Nominal Scales. Educational and Psychological Measurement. 1960;20:37–46.

22. Zhou Y, Zhou B, Pache L, Chang M, Khodabakhshi AH, Tanaseichuk O, Benner C, Chanda SK. Metascape provides a biologist-oriented resource for the analysis of systems-level datasets. Nat Commun. 2019;3:10–1523.

23. Thielens NM, Tedesco F, Bohlson SS, Gaboriaud C, Tenner AJ. C1q: A fresh look upon an old molecule. Molecular Immunology 2017;89:73–83.

24. Bulla R, Tripodo C, Rami D, Ling G-S, Agostinis C, Guarnotta C, Zorzet S, Durigutto P, Botto M, Tedesco F. C1q acts in the tumour microenvironment as a cancer-promoting factor independently of complement activation. Nature Communications 2016;7:10346.

25. Mangogna A, Belmonte B, Agostinis C, Zacchi P, Iacopino DG, Martorana A, Rodolico V, Bonazza D, Zanconati F, Kishore U, Bulla R. Prognostic Implications of the Complement Protein C1q in Gliomas. Frontiers in Immunology 2019;10:2366.

26. Zhang D, Li Y, Li H, Tang T, Zheng Y, Guo X, Xu X. A preliminary study of the complement component 1q levels in predicting the efficacy of combined immunotherapy in patients with lung cancer. Cancer Management Research 2021;13:7131–7.

27. Mangogna A, Agostinis C, Bonazza D, Belmonte B, Zacchi P, Zito G, Romano A, Zanconati F, Ricci G, Kishore U, Bulla R. Is the complement protein C1q a pro-or antitumorigenic factor? Bioinformatics analysis involving human carcinomas. Frontiers in Immunology 2019;10:865.

28. Kou W, Li B, Shi Y, Zhao Y, Yu Q, Zhuang J, Xu Y, Peng W. High complement protein C1q levels in pulmonary fibrosis and non-small cell lung cancer associated with poor prognosis. BMC Cancer 2022;22:110.

29. Kaur A, Sultan SHA, Murugaiah V, Pathan AA, Alhamlan FS, Karteris E, Kishore U. Human C1q induces apoptosis in an ovarian cancer cell line via tumour necrosis factor pathway. Frontiers in Immunology 2016;7:599.

30. Hong Q, Sze C-I, Lin S-R, Lee M-H, He R-Y, Schultz L, Chang J-Y, Chen S-J, Boackle RJ, Hsu L-J,. Chang N-S. Complement C1q activates tumour suppressor WWOX to induce apoptosis in prostate cancer cells. PLoS One 2009;4:e5755.

31. Agostinis C, Vidergar R, Belmonte B, Mangogna A, Amadio L, Geri P, Borelli V, Zanconati F, Tedesco F, Confalonieri M, Tripodo C, Kishore U, Bulla R. Complement protein C1q binds to hyaluronic acid in the malignant pleural mesothelioma microenvironment and promotes tumor growth. Frontiers in Immunology 2017;8:1559.

32. Zhang R, Liu Q, Li T, Liao Q, Zhao Y. Role of the complement system in the tumour microenvironment. Cancer Cell International 2019;19:300.

33. Cui J, Li G, Yin J, Li L, Tan Y, Wei H, Liu B, Deng L, Tang J, Chen Y, Yi L. GSTP1 and cancer: Expression, methylation, polymorphisms and signalling. International Journal of Oncology 2020;56:867–78.

34. Huang J, Tan P-H, Thiyagarajan J, Bay B-H. Prognostic significance of glutathione S-transferase-pi in invasive breast cancer. Modern Pathology 2003;16:558–65.

35. Meding S, Balluff B, Elsner M, Schöne C, Rauser S, Nitsche U, Maak M, Schäfer A, Hauck SM, Ueffing M, Langer R, Höfler H, Friess H, Rosenberg R, Walch A. Tissue-based proteomics reveals FXYD3, S100A11 and GSTM3 as novel markers for regional lymph node metastasis in colon cancer. Journal of Pathology 2012;228:459–470.

36. Cabelguenne A, Loriot MA, Stucker I, Blons H, Koum-Besson E, Brasnu D, Beaune P, Laccourreye O, Laurent-Puig P, De Waziers I. Glutathione-associated enzymes in head and neck squamous cell carcinoma and response to cisplatin-based neoadjuvant chemotherapy. International Journal of Cancer 2001;93:725–30.

37. Pectasides D, Kamposioras K, Papaxoinis G, Pectasides E. Chemotherapy for recurrent cervical cancer. Cancer Treatment Reviews 2008;34:603–13.

38. Checa-Rojas A, Delgadillo-Silva LF, Velasco-Herrera MdC, Andrade-Dominguez A, Gil J, Santillán O, Lozano L, Toledo-Leyva A, Ramirez-Torres A, Talamas-Rohana P, Encarnación-Guevara S. GSTM3 and GSTP1: novel players driving tumor progression in cervical cancer. Oncotarget 2018;9:21696–714.

39. Aitkenhead M, Wang S-J, Nakatsu MN, Mestas J, Heard C, Hughes CCW. Identification of endothelial cell genes expressed in an in vitro model of angiogenesis: induction of ESM-1, βig-h3, and NrCAM. Microvascular Research 2002;63:159–71.

40. He Y. Shao F, Pi W, Shi C, Chen Y, Gong D, Wang B., Cao Z, Tang K. Largescale transcriptomics analysis suggests over-expression of BGH3, MMP9 and PDIA3 in oral squamous cell carcinoma. PLoS One 2016;11:e0146530.

41. Srinivasan S, Guha M, Dong DW, Whelan KA, Ruthel G, Uchikado Y, Natsugoe S, Nakagawa H, Avadhani NG. Disruption of cytochrome c oxidase function induces Warburg effect and metabolic reprogramming. Oncogene 2016;35:1585–95.

42. Sánchez-Corral P, Criado García O, Rodríguez de Córdoba S. Isoforms of human C4b-binding protein. I. Molecular basis for the C4BP isoform pattern and its variations in human plasma. Journal of Immunology 1995;155:4030–6.

43. Sogawa K, Takano S, Iida F, Satoh M, Tsuchida S, Kawashima Y, Yoshitomi H, Sanda A, Kodera Y, Takizawa H, Mikata R, Ohtsuka M, Shimizu H, Miyazaki M, Yokosuka O, Nomura F. Identification of a novel serum biomarker for pancreatic cancer, C4b-binding protein α-chain (C4BPA) by quantitative proteomic analysis using tandem mass tags. British Journal of Cancer. 2016;115:949–956.

44. Pasic I, Shlien A, Durbin AD, Stavropoulos DJ, Baskin B, Ray PN, Novokmet A, Malkin D. Recurrent focal copy-number changes and loss of heterozygosity implicate two noncoding RNAs and one tumor suppressor gene at chromosome 3q13.31 in osteosarcoma. Cancer Research 2010;70:160–71.

45. Yen CC, Chen WM, Chen TH, Chen WYK, Chen PCH, Chiou HJ, Hung GY, Wu HTH, Wei CJ, Shiau CY, Wu YC, Chao TC, Tzeng CH, Chen PM, Lin CH, Chen YJ, Fletcher JA. Identification of chromosomal aberrations associated with disease progression and a novel 3q13.31 deletion involving LSAMP gene in osteosarcoma. International Journal of Oncology 2009;35:775–88.

46. Ntougkos E, Rush R, Scott D, Frankenberg T, Gabra H, Smyth JF, Sellar GC. The IgLON family in epithelial ovarian cancer: expression profiles and clinicopathologic correlates. Clinical Cancer Research 2005;11:5764–8.

47. Kanayama HO, Chen JD, Lui WO, Takahashi M, Kagawa S, Teh BT. LSAMP and norei: novel genes in hereditary and sporadic clear cell renal cell carcinomas (CCRCC). Journal of Urology 2004;171:200.

48. Chen J, Lui WO, Vos MD, Clark GJ, Takahashi M, Schoumans J, Khoo SK, Petillo D, Lavery T, Sugimura J, Astuti D, Zhang C, Kagawa S, Maher ER, Larsson C, Alberts AS, Kanayama HO, Teh BT. The t(1;3) breakpoint-spanning involved in clear cell renal cell genes LSAMP and NORE1 are carcinomas. Cancer Cell 2003;4:405–13.

49. Chang CY, Wu KL, Chang YY, Liu YW, Huang YC, Jian SF, Lin YS, Tsai PH, Hung JY, Tsai YM, Hsu YL. The Downregulation of LSAMP Expression Promotes Lung Cancer Progression and Is Associated with Poor Survival Prognosis. Journal of Personalised Medicine. 2021;11:578–592.

50. Kitazawa S, Takaoka Y, Ueda Y, Kitazawa R. Identification of calmodulin-like protein 5 as tumor-suppressor gene silenced during early stage of carcinogenesis in squamous cell carcinoma of uterine cervix. International Journal of Cancer. 2021;149:1358–1368.

51. Zheng T-L, Cao K, Lang C, Zhang K, Guo H-Z, Li D-P, Zhao S. Prognostic value of C-reactive protein in esophageal cancer: a meta-analysis. Asian Pacific Journal of Cancer Prevention 2014;15:8075–81.

52. Yadin DA, Robertson IB, McNaught-Davis J, Evans P, Stoddart D, Handford PA, Jensen SA, Redfield C. Structure of the fibrillin-1 N-terminal domains suggests that heparan sulfate regulates the early stages of microfibril assembly. Structure. 2013;21:1743–1756.

53. Kerslake R, Hall M, Vagnarelli P, Jeyaneethi J, Randeva HS, Pados G, Kyrou I, Karteris E. A pancancer overview of FBN1, asprosin and its cognate receptor OR4M1 with detailed expression profiling in ovarian cancer. Oncology Letters 2021;22:650.

54. Cierna Z, Mego M, Jurisica I, Machalekova K, Chovanec M, Miskovska V, Svetlovska D, Kalavska K, Rejlekova K, Kajo K, Mardiak J, Babal P. Fibrillin-1 (FBN-1) a new marker of germ cell neoplasia in situ. BMC Cancer 2016;16:597.

55. Cui F-L, Mahmud A-N, Xu Z-P, Wang Z-Y, Hu J-P. VASN promotes proliferation of prostate cancer through the YAP/TAZ axis. European Review for Medical and Pharmacological Sciences 2020;24:6589–6596.

56. Aydin M, Kızıltan R, Algul S, Kemik O. The utility of serum vasorin levels as a novel potential biomarker for early detection of colon cancer. Cureus 2022;14:e21653.

57. Zhang J, Shridhar R, Dai Q, Song J, Barlow SC, Yin L, Sloane BF, Miller FR, Meschonat C, Li BD, Abreo F, Keppler D. Cystatin m: a novel candidate tumor suppressor gene for breast cancer. Cancer Research. 2004;64:6957–64.

58. Shridhar, R., Zhang, J., Song, J. et al. Cystatin M suppresses the malignant phenotype of human MDA-MB-435S cells. Oncogene 2004;23:2206–2215.

59. Daniel K. Towards novel anti-cancer strategies based on cystatin function. Cancer Letters. 2006;235:159–176

60. Pérez-Holanda S, Blanco I, Menéndez M, Rodrigo L. Serum concentration of alpha-1 antitrypsin is significantly higher in colorectal cancer patients than in healthy controls. BMC Cancer 2014;14:355.

61. El-Akawi ZJ, Abu-awad AM, Khoun NA. Alpha-1 antitrypsin blood levels as indicator for the efficacy of cancer treatment. World Journal of Oncology 2013;4:83–6.

62. Short SP, Williams CS. Selenoproteins in tumorigenesis and cancer progression, Advances in Cancer Research 2017;136:49–83.

63. Takata Y, Kristal AR, Santella RM, King IB, Duggan DJ, Lampe JW, Rayman MP, Blount PL, Reid BJ, Vaughan TL, Peters U. Selenium, selenoenzymes, oxidative stress and risk of neoplastic progression from Barrett’s esophagus: results from biomarkers and genetic variants. PLoS One. 2012;7:e38612.

64. Maher SG, McDowell DT, Collins BC, Muldoon C, Gallagher WM, Reynolds JV. Serum proteomic profiling reveals that pretreatment complement protein levels are predictive of esophageal cancer patient response to neoadjuvant chemoradiation. Ann. Surg. 2011;254:809–816.

