## Supplementary data for "Identification of plasma proteins associated with oesophageal cancer chemotherapeutic treatment outcomes using SWATH-MS"

**Table S1. Proteins exhibit disagreement between depleted and non-depleted plasma samples.**

**Figure S1. Median normalisation on protein level for empirical bayes analysis.**

**Figure S2. Median normalisation on peptide level for empirical bayes analysis.**

**Figure S3. Comparison of peptide level data from depleted vs non-depleted plasma samples.**

**Figure S4. Empirical Bayes result on relative protein abundance in non-depleted plasma.**

**Figure S5. Multivariate predictive modelling on peptide relative intensities for proteins with multiple peptides identified in non-depleted data.**

**Figure S6. Relative intensities of peptides predictors for proteins that can classify patient samples into two TRG groups (Average Cohen's kappa > 0.35, ranked by average classification accuracy).**

**Table S1. Proteins that exhibit disagreement between depleted and non-depleted plasma samples.**

| **Protein** | **No. peptide in depleted sample** | **No. peptide in non-depleted sample** | **No. shared peptide** | **Difference (non-depleted - depleted)** |
| --- | --- | --- | --- | --- |
| ABCA1 | 1 | 1 | 1 | >1.96SD |
| ALBU | 3 | 4 | 3 | >1.96SD |
| IGHG4 | 1 | 6 | 1 | >1.96SD |
| IGHG2 | 2 | 4 | 2 | >1.96SD |
| IGHG3 | 1 | 5 | 1 | >1.96SD |
| DSG1 | 9 | 1 | 1 | <-1.96SD |
| TFR1 | 14 | 1 | 1 | <-1.96SD |
| CATA | 9 | 2 | 2 | <-1.96SD |
| FAM3C | 2 | 1 | 1 | <-1.96SD |
| BIG3 | 2 | 1 | 1 | <-1.96SD |
| MENT | 1 | 1 | 1 | <-1.96SD |
| TALDO | 2 | 1 | 1 | <-1.96SD |
| CO6A1 | 3 | 1 | 1 | <-1.96SD |
| LDHB | 6 | 1 | 1 | <-1.96SD |
| BGH3 | 13 | 1 | 1 | <-1.96SD |
| IBP6 | 2 | 1 | 1 | <-1.96SD |

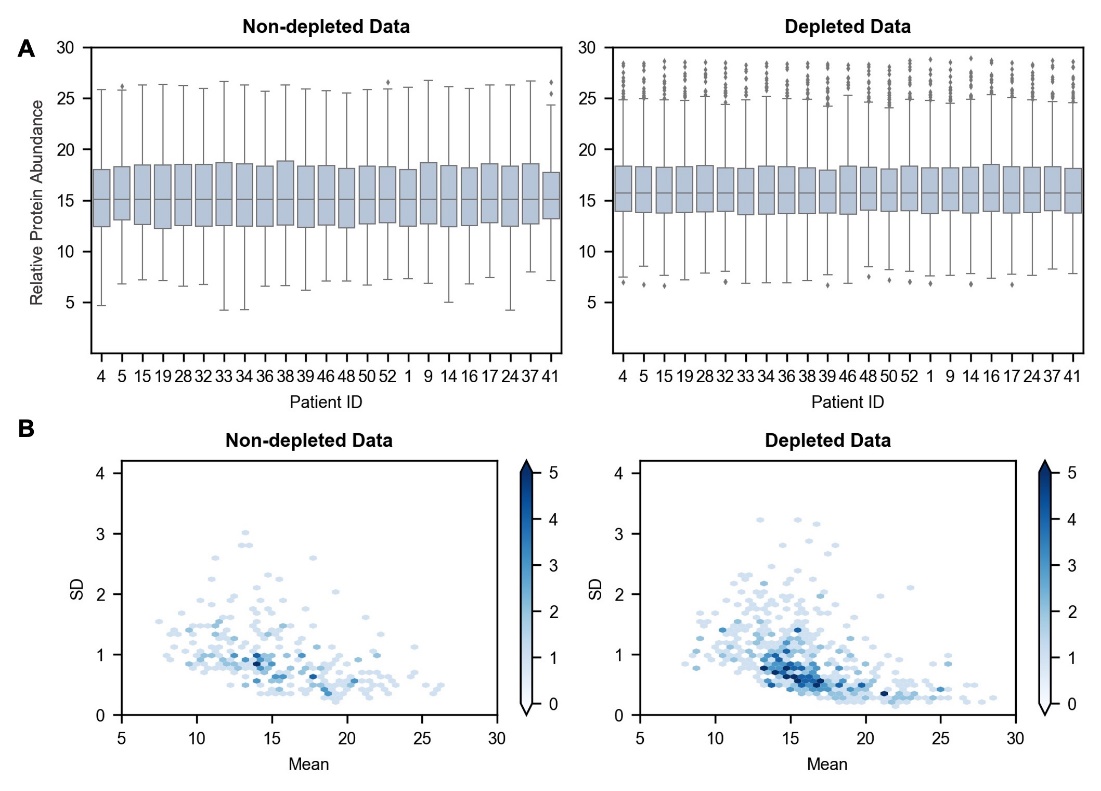

**Figure S1. Median normalisation on protein level for empirical bayes analysis.** (A). Boxplots of Relative protein intensity for each patient. After log transformation and median normalisation, all samples have an equal median of relative protein abundance. (B). The mean-variance plots for normalised protein intensities for depleted data and non-depleted data respectively. Normalisation effectively stabilised variance across the mean intensities of peptides and scaled the patient samples to the same median.

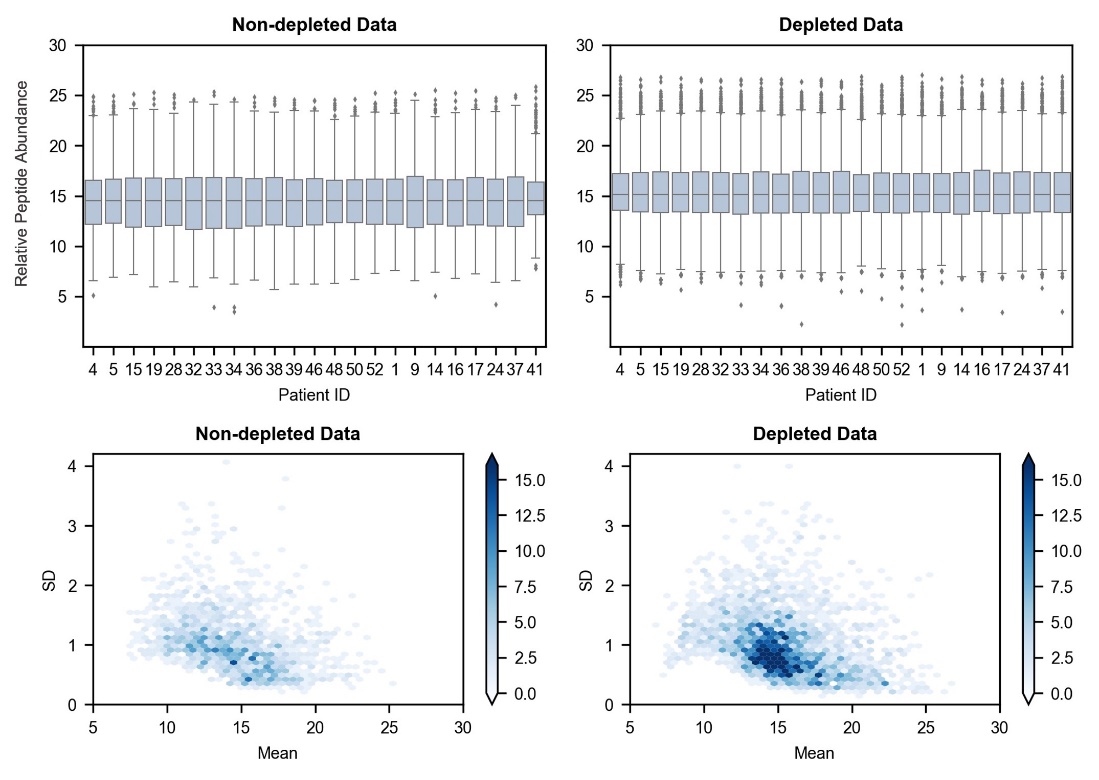

**Figure S2. Median normalisation on peptide level for empirical bayes analysis.**  (A). Boxplots of Relative protein intensity for each patient. After log transformation and median normalisation, all samples have an equal median of relative peptide abundance. (B). Median normalisation on peptide level for multivariate LDA model. The mean-variance plots for normalised peptide intensities for depleted data and non-depleted data are presented. Normalisation effectively stabilised variance across the mean intensities of peptides and scaled the patient samples to the same median.

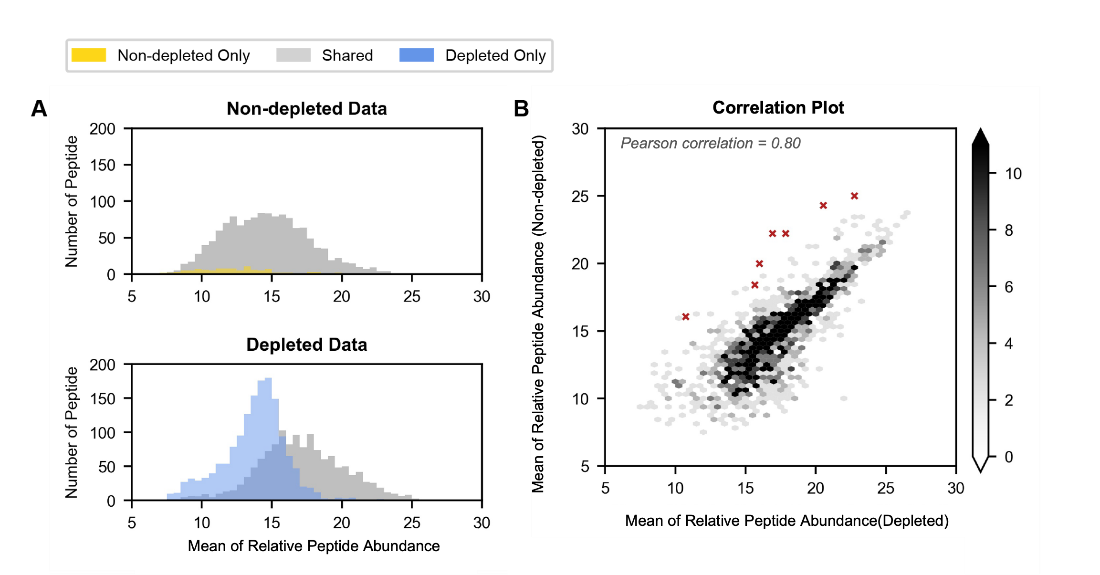

**Figure S3. Comparison of peptide level data from depleted vs non-depleted plasma samples.** A. Distributions of means of commonly found peptides from non-depleted data and depleted data. B. the scatterplot coloured by density showing the correlation between depleted data and non-depleted data for all common peptides. Note that albumin and immunoglobin peptides (red crosses) have a higher abundance in non-depleted data than in depleted data as expected.

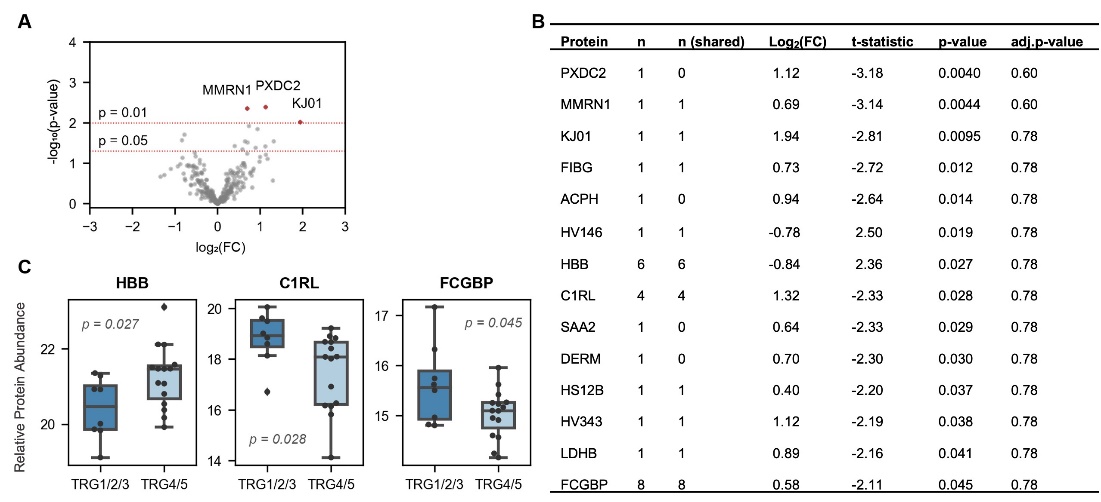

**Figure S4. Empirical Bayes result on relative protein expressions in non-depleted plasma.** A. Volcano plot showing result of non-depleted plasma. It is based on log_2_-fold change (x-axis) against negative log_10_ p-value (y-axis) and orange dots represent proteins with p-value<0.01 identified by empirical bayes test. B. Quantitative data for proteins with p-value<0.05. n: number of peptides identified from non-depleted plasma sample; n (shared): number of peptides that are commonly found in depleted plasma sample. C. Boxplots showing the relative expression of differentially expressed proteins in TRG1/2/3 and TRG4/5 group (only quantification is based on more than one peptide is shown).

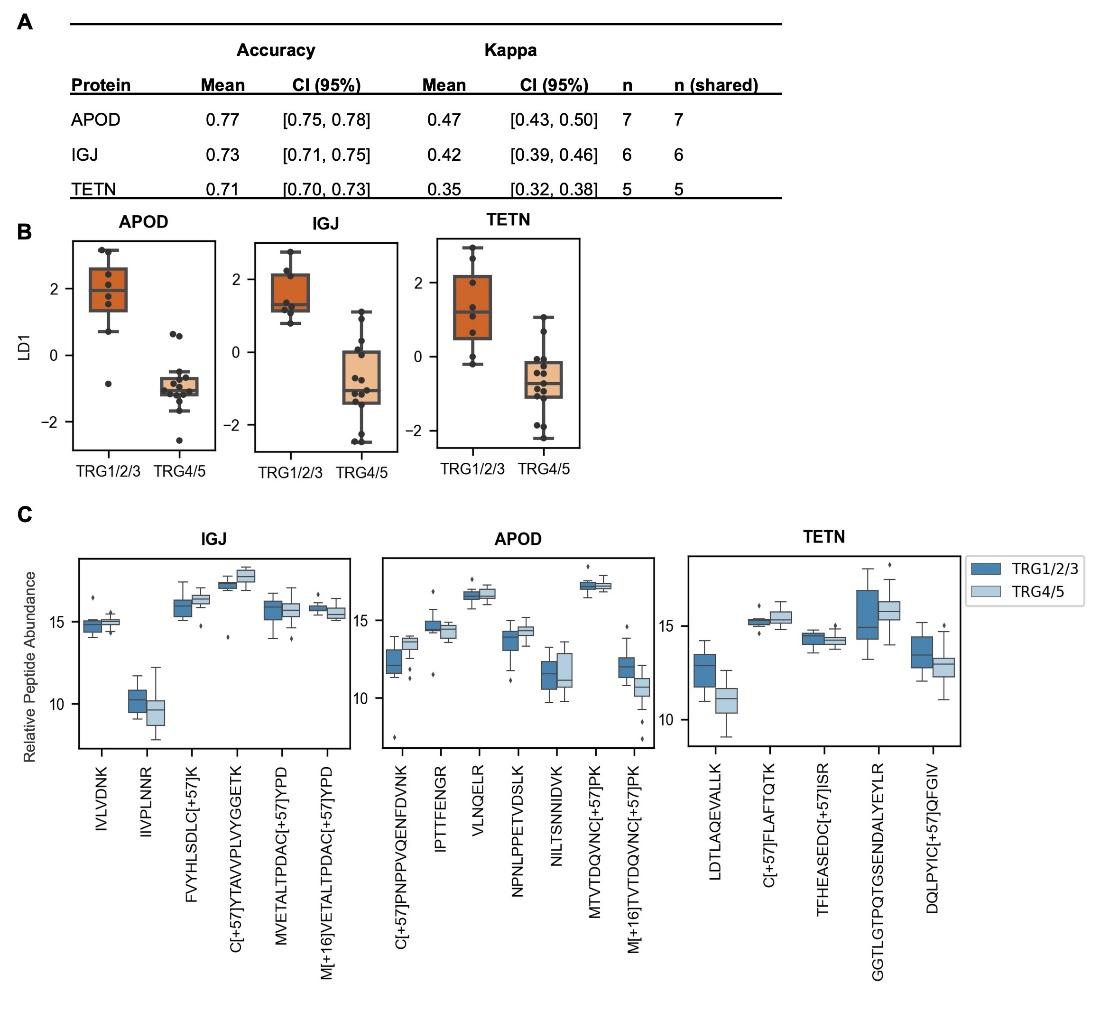

**Figure S5. Multivariate predictive modelling on peptide relative intensities for proteins with multiple peptides identified in non-depleted data.** A. Predictive performance for proteins with multiple peptides in non-depleted data. Proteins are selected and ranked by the average of Cohen's kappa (kappa: mean > 0.35) and classification accuracy, respectively. CI: confidence interval. n: number of peptides identified from depleted plasma sample; n (shared): number of peptides that are commonly found in non-depleted plasma samples). B. Linear discriminant scores regarding low (1/2/3) and high (4/5) TRG scores. Significant differences along the first linear discriminant LD1 were detected for all the selected proteins, indicating the classification ability of each protein based on the weighted sum of peptides. C. Relative expression of peptides as predictors in each LDA model. Relative peptides intensities are multiplied with the corresponding coefficients and summed to form the decision rule for each classification task. Patient samples are classified based on the weighted sum values, and the coefficient indicates the contribution of each peptide predictor. The LDA model has the advantage to detect peptides that show contrast relative expressions in low (1/2/3) and high (4/5) TRG score groups, which has been cancelled out at protein relative expression level and thus not discovered by univariate t-test. As shown in the figure, IGJ, APOD and TETN proteins have peptides showing both higher and lower relative expressions in the TRG1/2/3 group compared to the TRG4/5 group.

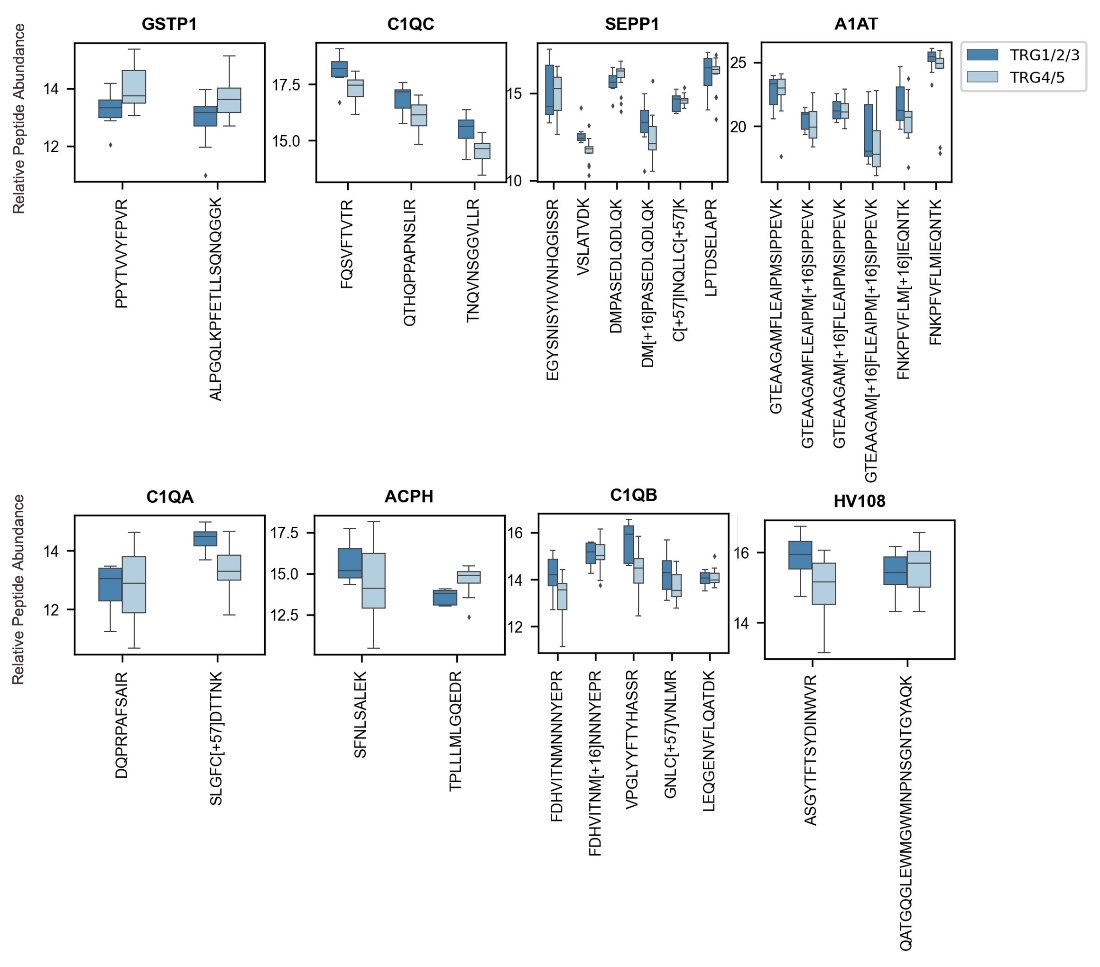

**Figure S6. Relative intensities of peptides predictors for proteins that can classify patient samples into two TRG groups (Average Cohen's kappa > 0.35, ranked by average classification accuracy).** Relative peptide abundance is multiplied with the corresponding coefficients and summed to form the decision rule for each classification task. Patient samples are classified based on the weighted sum values, and the coefficient indicates the contribution of each peptide predictor. The LDA model has the benefit of identifying proteins that have peptides that show contrast relative expressions in low (1/2/3) and high (4/5) TRG score group (e.g., in SEPP1, ACPH and HV108 proteins, there are peptides showing both higher and lower relative abundance in the TRG1/2/3 group compared to the TRG4/5 group, thus has been cancelled out at protein relative abundance level), and the ones that did not reach the level of significance at relative protein intensities (e.g., C1QA, A1AT) by taking advantage of the correlation between peptides.
